# Updated reference genome sequence and annotation of *Mycobacterium bovis* AF2122/97

**DOI:** 10.1101/092122

**Authors:** Kerri M. Malone, Damien Farrell, Tod P. Stuber, Olga T. Schubert, Reudi Aebersold, Suelee Robbe-Austerman, Stephen V. Gordon

## Abstract

We report an update to the reference genome of the bovine tuberculosis bacillus *Mycobacterium bovis* AF2122/97 generated using an integrative multi-‘omics approach. Updates include 42 new CDS, 14 modified annotations, 26 SNP corrections, and disclosure that the RD900 locus, previously described as absent from the genome, is in fact present.

## Text

The *Mycobacterium tuberculosis* complex (MTBC) is a collection of genetically related mycobacterial species that cause tuberculosis (TB) in human and animal hosts. *Mycobacterium bovis*, the causative agent of bovine tuberculosis (bTB), is the most widely studied animal-adapted MTBC member; bTB exacts a tremendous global economic toll through productivity losses and disease control costs, while zoonotic transmission of *M. bovis* infection is a threat to human health (Abernethy *et al*., 2013; De Garine-Wichatitsky *et al*., 2013; Kazoora et al., 2016; Khattak *et al*., 2016; Malama *et al*., 2014; Muller *et al*., 2013).

*M. bovis* AF2122/97 was the first bovine MTBC strain to be fully sequenced and provided a reference genome (Garnier et al., 2003). Initial comparisons of the *M. bovis* AF2122/97 genome with that of the human-adapted *M. tuberculosis* H37Rv reference genome revealed high nucleotide sequence identity (> 99%), no unique genes *per se* in *M. bovis* AF2122/97 and a number of genomic deletions that led to a reduced genome size (Garnier et al., 2003). *M. bovis* AF2122/97 continues to serve as an MTBC reference genome despite last being updated in 2003; by comparison, the genome annotation of the reference *M. tuberculosis* H37Rv strain is currently on release 27 (Lew *et al*., 2013). An updated reference *M. bovis* genome will provide an essential resource for the TB research community and as a basis for comparative studies into animal- and human-adapted MTBC members.

To update the *M. bovis* AF2122/97 genome and annotation, a low-passage stock taken from the original *M. bovis* AF2122/97 seed stock was re-sequenced and re-annotated using a combination of DNA-, RNA-sequencing and proteomics data. All nucleic acid and protein samples were derived from exponentially grown cultures.

Short read DNA sequencing libraries were prepared using the Nextera XT DNA Library Preparation Kit (Illumina^®^) and sequenced on the MiSeq^®^ system (Illumina^®^), generating 250bp paired-end reads that were trimmed using Sickle (Q >30), with 60X reference coverage (Joshi NA, 2011). For PacBio RS II sequencing, enzymatically extracted DNA was prepared using large insert library (6kb-8kb) size selection (van Soolingen *et al.,* 1991). Two SMRT cells were used for an output of 542,585,804 bases, a mean read length of 8,141, and 86X reference coverage. DNA sequencing datasets were analysed using a combination of *de novo* assembly (short reads, SOAPdenovo (Xie *et al.,* 2014); long reads, Canu (Koren S, 2016)) and nucleotide variant identification methods (short reads, Stampy, SAMtools and VCFtools (Li, 2011; Li *et al*., 2009; Lunter and Goodson, 2011); long reads, Pilon (Walker *et al*., 2014); MUMmer (Kurtz *et al*., 2004)). This allowed for the update of the genome nucleotide sequence and the identification of genomic regions that were misassembled, or missed entirely, in the original sequencing project. Re-annotation of the *M. bovis* AF2122/97 genome was achieved by automatic annotation transfer from *M. tuberculosis* H37Rv (Version 27) (Otto et al., 2011) and a proteogenomic analysis using both M. bovis AF2122/97 shotgun MS/MS, SWATH MS datasets and *M. tuberculosis* H37Rv SWATH MS datasets (Schubert et al., 2013).

Overall, 26 single nucleotide polymorphisms were identified. Strikingly, the large sequence polymorphism RD900, originally described as deleted from *M. bovis* 2122/97 (Bentley *et al*., 2012), was found to be present; recombination between repeat structures flanking the RD900 locus in clones used for the original shotgun sequencing genome project may have led to loss of RD900. Furthermore, 42 novel coding sequences were identified while 14 existing annotations were modified.

**Nucleotide sequence accession number(s):** This Whole Genome Shotgun project had been deposited in DDBj/ENA/Genbank under the accession no. LT708304. SWATH MS data can be found on PeptideAtlas (http://www.peptideatlas.org) under identifier: PASS00932.

## Acknowledgements

The work was supported by funding from the Department for Agriculture, Food and the Marine (MycobactDiagnosis 11/RD/EMIDA/1), Science Foundation Ireland (08/IN.1/B2038), SystemsX.ch (through the project TbX) and a research grant from Institut Mérieux. We would like to thank Dr. Gerard Cagney (UCD School of Biomolecular and Biomedical Science) and Dr. Kieran Wynne (UCD Conway Proteomics Core) for shotgun mass spectrometric analysis of *M. bovis* samples.

